## Supplemental information for "Diversification of *Pseudomonas aeruginosa* biofilm populations under repeated phage exposures decreases the efficacy of the treatment"

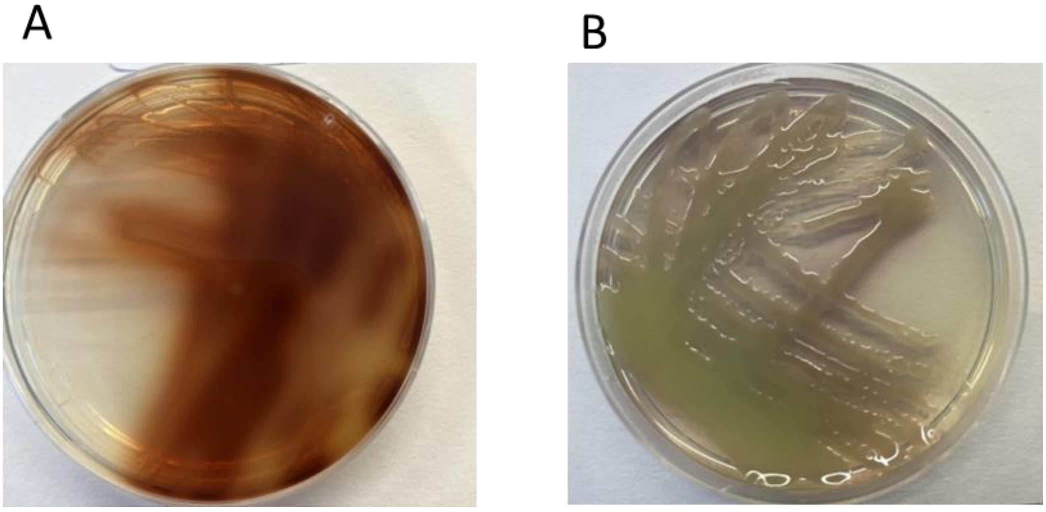


**Figure S1.** Representative pictures of *P. aeruginosa* colonies producing pyomelanin (A) and alginate (mucoid colony) after bacteriophage treatment of biofilm.

### Supplementary Results

#### Additional genetic alterations in clones collected from biofilm exposed to phages

Upon comparing phage-treated clones with their ancestors, additional alterations were discerned that do not exhibit a direct correlation with the phage susceptibility phenotype (Figure 5 A to E). The predominant mutation identified in five resistant and three sensitive clones of the clinical isolate CF341_08 was the in-frame insertion Leu486insGluPro in the *dnaX* gene encoding for the DNA polymerase III subunits γ and τ. The DNA polymerase III is the major conserved DNA replication holoenzyme in prokaryotes. The mutation results in the extension of the GluPro compositionally biased (CB) region, consisting of 13 GluPro repeats, by one additional repeat. The consequences of this alteration remain uncertain, although it appears more probable that it represents a common mutation within this gene, given its occurrence in a highly repetitive sequence.

In one NP3-resistant clone of PA14, a synonymous mutation manifested in the *ubiA* gene encoding for 4-hydroxybenzoate octaprenyltransferase, the second enzyme in the pathway of ubiquinone biosynthesis. This mutation does not impart any consequential alterations to the protein's functionality, but potential impacts related to codon usage leading to a diminished ubiquinone content cannot be entirely ruled out. In general, these mutations surfaced early in the biofilm population (after 24 h), and were unique to each strain lineage, underscoring their spontaneous nature.

The majority of PAO1 subjected to NP3 and M32 treatment exhibited a Ser125Gly substitution in RluA and a frameshift at Thr46 in Wzy, independently of treatment duration or phage susceptibility phenotype. The RluA modifies primary the U-nucleotides in the tRNA[1]. The Ser125Gly substitution in RluA occurred between two β-sheets. A *blastp* analysis revealed the presence of Gly at this position in RluA proteins from other *P. aeruginosa* strains, suggesting that this alteration is likely inconsequential to the protein's function.

Various frameshifts or nonsense mutations, all resulting in the loss of function, were found in PilY1 in both in M32-sensitive and M32-resistant clones, as well as in one NP3-resistant clone of PAO1. PilY1 is a component of the inner membrane sub-complex of the type IV pilus assembly machinery [2]. As PilY1 is situated internally, it is improbable that it plays a role in phage recognition, as it is not exposed to the external environment. Malfunctions in this protein do not seem to induce alterations in other recognition patterns associated with pili that typically accompany phage resistance.

The predominant mutation observed early in the biofilm population of CF341_06, present in both phage-sensitive and -resistant clones, involved a duplication of Pro at amino acid position

289 in the type VI secretion system (T6SS) forkhead-associated protein Fha1. This duplication occurred in the C-terminal end of the distorted loop, not in the forkhead-associated domain. Fha1 is located in the inner membrane at the T6SS-membrane core complex and activates pilus assembly upon phosphorylation, triggered by perturbations of the outer membrane [3]. Given that the Pro duplication occurs at the termination of the GlnPro compositional bias region (consisting of 11 GlnPro repeats), a malfunction of Fha1 and subsequent absence of T6SS pili is deemed rather unlikely, although it cannot be completely excluded. Importantly, Fha1 is not accessible to the phage as a receptor, making it also improbable that this alteration impacts host recognition by the NP3 phage.

**Table S2. Information on CF clinical isolates (CF 341)**

| Patient CF 341 | | |
|---|---|---|
| Isolate | CF341_06_NM | CF341_08_NM |
| Year of isolation | 2006 | 2008 |
| Nonmucoid (N) phenotype | NM | NM |
| Infection status at time of isolation | Intermittently colonised | Chronically infected |
| *P. aeruginosa* precipitin | 1 | 28 |
| Weeks of peroral/inhalation/ intravenous antibiotic therapy | 0 | Po: 6<br>IV: 14 |
| Sensitivity towards ciprofloxacin of the bacterial isolates in planktonic form (µg/mL) determined by e-test | 0.064 | 0.380 |
| Generation time *in vitro* (min.) | 42.2 | 47 |

**Table S3. Host-range NP3 determined by spot assay on environmental, reference and CF clinical isolates.**

| Source | *P. aeruginosa* strain | Lysis by NP3 |
|---|---|---|
| environmental | PS F8 | + |
| environmental | PS F10 | + |
| environmental | PS 119X | + |
| environmental | PS M4 | + |
| reference | PAO1 | + |
| reference | PDO300 (M) | + |
| reference | PA14 | + |
| clinical | CF 46 476a/88 | + |
| clinical | CF 46 476b/88 | + |
| clinical | CF46 52329a/97 | + |
| clinical | CF 46 52329b/97 | + |
| clinical | CF46 1395a/03 | + |
| clinical | CF 46 1395b/03 | - |
| clinical | CF 30 19731a/92 | + |
| clinical | CF30 19731b/92 | + |
| clinical | CF30 75887a/01 | + |
| clinical | CF30 75887b/01 | + |
| clinical | CF128 1398a/92 | + |
| clinical | CF128 1398b/92 | + |
| clinical | CF128 8503a/02 | + |
| clinical | CF128 8503b/02 | 0 |
| clinical | CF 89 15061/78 | + |
| clinical | CF238 82847b/01 | + |

| | | |
|---|---|---|
| clinical | CF 341_ 28898_06 | + |
| clinical | CF 341_43885A 08A (M) | (+m) |
| clinical | CF 341 43885B_08 (NM) | + |
| clinical | CF 405_56281A_05 | + |
| clinical | CF 405_3299_10 | + |
| clinical | CF 408_19970_06 | + |
| clinical | CF 408_38574A_06 | - |
| clinical | CF 414_15321_06 | + |
| clinical | CF 414_18275_08 | + |
| clinical | CF 414_36687_09 | - |
| clinical | CF 431_24293_08 | + |
| clinical | CF 431_7494_11 | + |
| clinical | CF 483_7161A_04 | + |
| clinical | CF 483_2728A_06 | (+) |
| clinical | CF 483_2728B_06 | (+) |
| clinical | CF 519_19637B_07 | + |
| clinical | CF 519_17006_1_11 | + |
| clinical | CF 519_17006_2_11 | + |
| clinical | CF 544_55286B_10 | + |
| clinical | CF 544_60163B_11 | + |
| clinical | CF 544_60163A_11 | + |
| clinical | CF 312_16704_91 | + |
| clinical | CF 312_20441A_95 | + |
| clinical | CF 312_20441B_95 | + |
